# Design and development of potent h-ACE2 derived peptide mimetics in SARS-CoV-2 Omicron variant therapeutics

**DOI:** 10.1101/2022.02.01.478632

**Authors:** M. L. Stanly Paul, Swathi Nadendla, M Elizabeth Sobhia

## Abstract

The pandemic of COVID-19 has become the global health challenge due to the emergence of new variants. The Receptor binding domain (RBD) of spike protein that makes direct interaction with ACE-2 has shown unique mutated residues in most of the variants of concern (VOC). Recently WHO declared the Omicron (B.1.1.529) as VOC considering it as a highly mutated variant which includes a total of 60 mutations out of which 15 mutations occurred in RBD region of SARS-CoV-2. Inhibition of Protein-protein (Omicron RBD-h-ACE2) interaction was already proved to inhibit the viral infection. In this study, by using molecular dynamic simulations efforts are made to explore the atomistic details of Omicron RBD-h-ACE2 interaction. Based on MD simulations, h-ACE2 motif is found to be interacting with omicron RBD domain. Interaction analysis had provided key residues interacting with Omicron-RBD that helped to extract h-ACE2 peptide. Here, rational design of the peptides that have resemblance with h-ACE2 is done and the peptide library is subjected for inhibition studies against Omicron-RBD. The current study helped to identify the significant peptides that can inhibit Omicron-RBD. Altogether the performed studies will provide an opportunity to develop potential therapeutic peptidomimetics effective against Omicron variant of SARS-CoV-2.

## INTRODUCTION

Since its emergence of SARS-CoV-2 in Wuhan province in China (2019) the impacts created by SARS-CoV-2 and its variants are having impeccable effect on the society and health care system worldwide^1^. As of 31^st^ January 2022, SARS-Cov-2 majorly Omicron variant has led to largest surge of conformed cases (375,241,633) with high mortality (5,681,741) rate in the world ^2^. In the last quarter of the year-2021 with the help of genomic sequencing first SARS-Cov-2 Omicron variant (B.1.1.529) cases were identified in South Africa. Recently, WHO classified the new variant Omicron (B.1.1.529) as Variant of Concern due to its higher infectivity rate.^3^. Omicron is considered as the most heavily mutated variant that has emerged so far with enhanced transmissibility and partial resistance to vaccine induced immunity^4, 5^. However, its closer sequence identity (99.63%) with alpha (B.1.1.7) variant suggests that omicron primary emergence in the population is way beyond its expectation even in vaccinated population.

Genomic studies on the variant revealed that there are fifteen mutations in the omicron RBD region out of which five mutations are present at the core binding region. Three mutations S371L, S373P, S375F in the subdomain are clustered at a hairpin loop that has initiated a main conformational change compared to other mutated RBD structures in other variants. Studies revealed that the interacting residues are same when compared to wild type RBD indicating that the overall binding mode is conserved^6^.

Spike protein is not only responsible for interaction with ACE-2 through RBD for entry into the host cell but is also a major inducer of neutralizing antibodies^7^. SARS Cov-2 is actively acquiring resistance since its emergence, hence cocktail of antibodies are required to produce desired clinical response whose production is comparatively expensive^8^. Inhibiting RBD-ACE-2 interactions by using peptides and peptidomimetics offers various advantages like high specificity and affinity and their ability to penetrate cell membranes^9^. Here, efforts are made for the design of peptide mimetic inhibitors of SARS-CoV-2 by mutation that are effective against the omicron variant^10^.

## MATERIALS AND METHODS

For this study the, new SARS-CoV-2 variant genome sequence was taken from the GISAID Database (GISAID accession EPI_ISL_6752027) which was submitted on November 22, 2021 by a team of researchers from Botswana-Harvard HIV Reference laboratory using a Nanopore MinION device^11^. As a part of structure prediction of omicron RBD, the template selection was done by using BLAST-P search. Native crystal structure of RBD-ACE2 alpha variant complex in PDB (7NXA)^12^ was taken as template among all others by considering the highest sequence identity of 93.30%. The multichain (omicron-RBD with ACE2) model structures generated through Modeller are further subjected to structure refinement.

### Comparative Electrostatic models

Electrostatic potential analysis provides guided parameters to design specific inhibitors against any biomolecules. But classical electrostatic potential analysis does not provide precise parameters due to non-consideration of solvent medium. Gaussian based smooth dielectric function considers the solvent medium that provides accurate computational calculations of electrostatic potentials. Here, Gaussian DelPhi program was used to analyze the electrostatic potential of both WT-RBD and omicron-RBD.

### MD simulations

Both WT-RBD and omicron-RBD along with hACE2 was subjected to 400ns of molecular dynamics simulation study to check folding and structural stability^13^. Desmond-maestro is used to prepare the complexes by adding TIP3P water, and neutralize the complexes by adding counter Na+ ions. Periodic boundary conditions (PBC) and OPLS3e forcefields were applied on the system before subjecting to the minimization process. Here we used Desmond standard NVT ensemble where constant pressure(1 bar) and brendsen thermostat with constant(300K) is maintained. Hydrogen bond restrains were considered by adding SHAKE algorithm whereas electro static forces by Particle Mesh Ewald (PME) algorithm in the MD run. Both WT and Omicron RBD-hACE2 systems were subjected to 400ns MD run.

### Designing peptide mimetic library

Most of the interactions between RBD-ACE2 are found to be from the α-1 helix of hACE2. So, mutating the residues of this helix may result in peptides that have higher binding affinity than ACE-2 becomes helpful in discovering of novel peptide inhibitors that prevent the RBD-ACE-2 interaction^10^. Here, the reference alpha helix peptide was taken from the ACE2-RBD homology model complex. The parent ACE2 alpha helix peptide in between Ile21 to Ser44 i.e. IEEQAKTFLDKFNHEAEDLFYQSS was found to interact with key binding residues of Omicron RBD. Based on the properties of amino acids on the apopeptide (*Figure* 1) design of peptide mutant library (456 peptidomimetics) over omicron RBD is done by carefully considering the structural changes of the protein, for this process in-house script was used. The generated peptidomimetic library was systematically divided based on the properties shown on helix wheel (Figure 1F).

**Figure 1:**
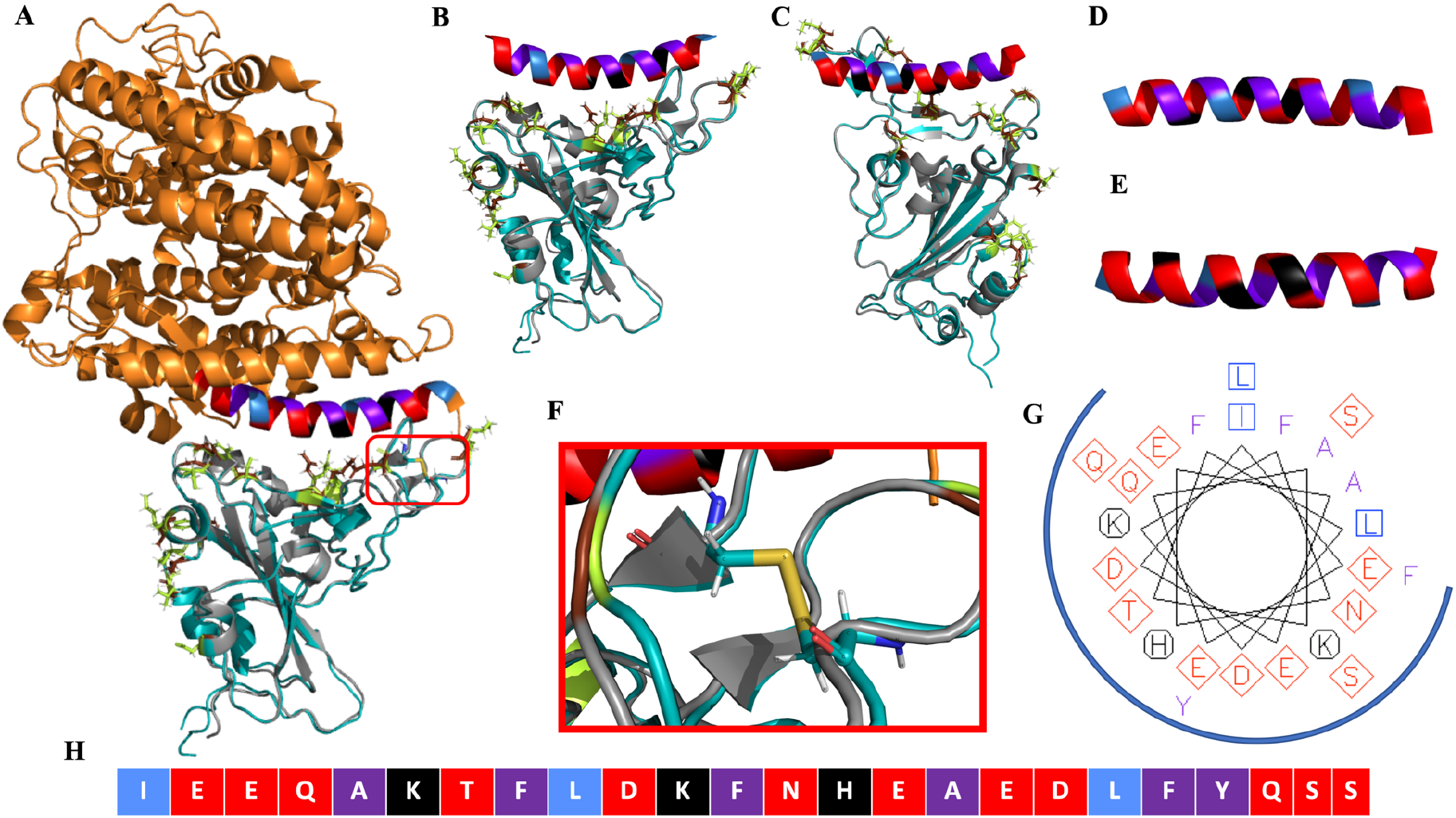
Cyan (omicron-RBD) Orange (ACE2) Gray(6LZG-RBD) Purple blue (mutated omicron RBD residues) Lime (Wildtype residues) **(A)** Superimposition of h-ACE2-omicron-RBD with wild type RBD **(B)** front view of RBD with peptide **( C)** back-side view of RBD with peptide **(D)** non-interacting surface of peptide **( E)** interacting surface of peptide **(F)** Disulphide bond between CYS480-CYS488 (**G)** Helical wheel representation of properties of peptide (diamond/red:- (DE negatively charged amino acids) and (NQST uncharged amino acids) negatively charged amino acids; octagon/black:- (HKR amino acids) positively charged amino acids; Squares:- (ILVM amino acids)aliphatic residues; no wheel:- (Y amino acid); amphipathic:- square (IL amino acids); Blue ring represents the interacting surface of h-ACE2 peptide **(H)** Colour Representation of amino acids according to properties (Red:-polar/negatively charged Black:- positively charged)

Initially, a 453 membered peptide mimetic library was modelled by using inhouse scripts and subjected to molecular docking over Omicron RBD binding site.

### Molecular docking of peptides

High ambiguity driven protein-protein docking (HADDOCK) is proved to be best docking server for protein-protein and protein-peptide docking. One feature that extricates it from other accessible software’s is the usage of experimentally derived data from NMR, mutagenesis and the provided solution was found to be identical with experimentally derived structure. Both the interacting residues in peptide and protein are given as input along with the active site residues of receptor. By defining Ambiguous interacting residues (AIRS) the main advantage is that only the poses in which the ligand interacts with the active residues are sampled rather than selection based on conformational energies. A three step protocol is used for both the proteinprotein and protein-peptide docking where step-1 contains rigid docking process, step-2 contains semi flexible docking and step-3 includes refining structures in presence of water. Docking score is mainly dependent on the summation of reports got through these three steps (HADDOCK docking score = 1.0EvdW + 0.2Eelec + 1.0Edesol + 0.1EAIR)^10^. A detail docking cluster reports were generated by HADDOCK server. A default cut-off of RMSD (2Å) is set and acceptable structures are ranked among the clusters.

Screening of peptide mimetics on Omicron RBD has provided key information on binding residues^10^. Out of the plethora of peptide library, fourteen peptides showed higher inhibition effect when compared with reference α-1 helix. Overall, the studies allowed to design novel peptide and peptidomimetic inhibitors that competitively bind to omicron RBD.

## RESULTS

The omicron RBD-ACE2 complex structure is modelled by using multichain script of Modeller. Initial protein interaction analysis provided binding residues with buried SASA(TYR489, TYR501, GLY502, SER496, PHE456, ARG498, ALA475, THR500, HID505, GLY476, ASN487, ARG493, TYR-453, SER494, TYR449) on RBD; Salt bridges ARG493-GLU35. П-П stacking(TYR501-TYR41) and vdW interactions were observed in between the following residues (PHE456-LYS31, THR500-TYR41, GLY476-GLN24, ARG493-HID43, SER494-HID34). Comparison of wildtype and omicron-RBD binding site revealed that omicrons mutated residues are noticeable longer side chains.

Comparative analysis of electrostatic surface of omicron-RBD with wild type RBD by using gaussian based dielectric function has given structural insights in the interaction site^14^. The electrostatic potential analysis of omicron-RBD has shown significant variation in charge potential, where in omicron-RBD, highly positive charged surface is observed when compared to the wild type.

**Figure 2:**
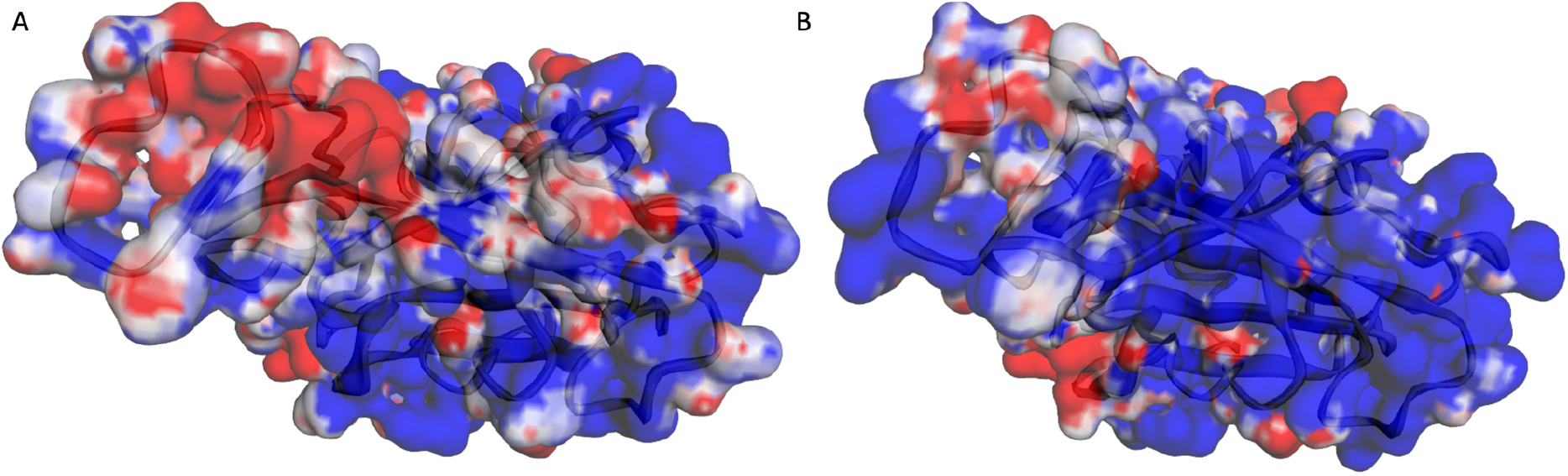
Red(Positive) and Blue(negative) colours denotes charge potential **(A)** Gaussian based smooth electrostatic potential of h-RBD **(B)** Gaussian based smooth electrostatic potential of Omicron-RBD.

This elucidates the significance of mutations in the tight binding process of omicron-RBD towards overall negatively charged h-ACE2. The modelled omicron-RBD in complex with ACE-2 is subjected to molecular dynamic simulations to get atomistic details of protein-protein interaction.

### Molecular dynamics simulation analysis

Time dependent all atom molecular dynamics simulations is already proved to be an efficient tool to analyse the folding behaviour of the protein in biological conditions. During whole production run RMSD and RMSF is useful to investigate the structural changes and to assess the stability the protein complex. In our study, initially RMSD of C-alpha-backbone of Omicron-RBD-h-ACE2 complex is raised from 2.4 to 3.7Å, but in omicron-RBD RMSD raised from 2.7 to 3.3Å over the time period of 20ns whereas in h-ACE2 RMSD raised from 2.6 to 3.2Å over the time period of 70ns and remained stable over the period of 400ns. In the process of Omicron RBD stabilization, clear structural changes were observed stepwise over the time period of 400ns; step-1 is observed in between 20ns to 220ns at 2.7 to 3.3 Å and step-2 from 320 to 400ns 3.7 to 4.5 Å where the structure got stabilized (Figure 3).

**Figure 3:**
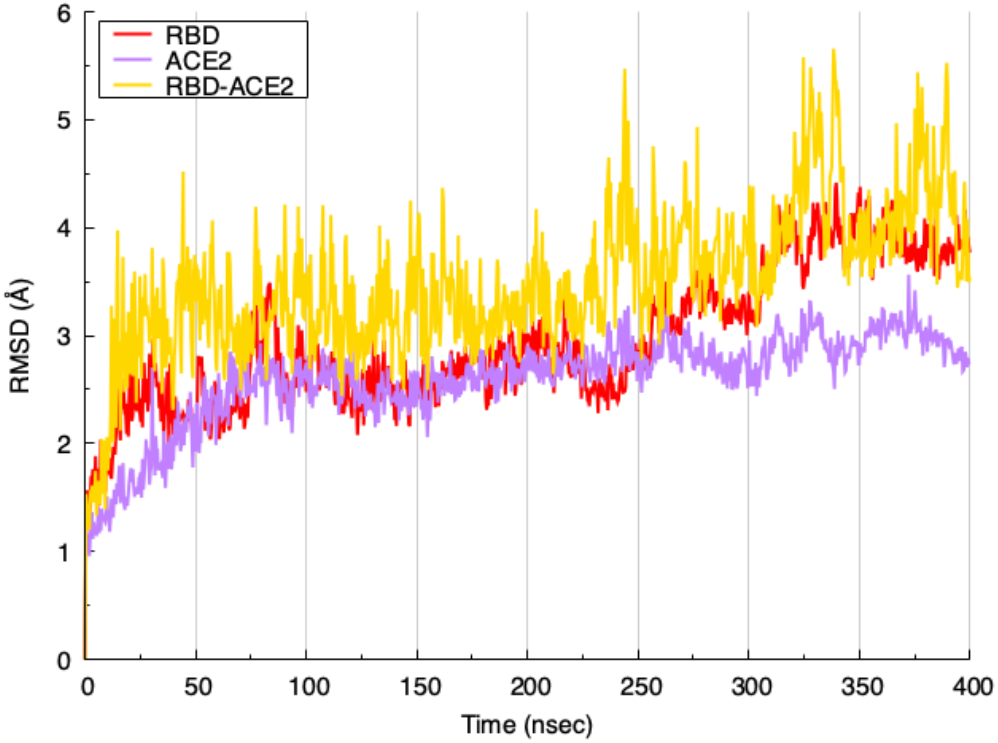
RMSD of omicron-RBD and h-ACE2 in complex(Yellow), Human-ACE2 (Purple) Omicron-RBD(Red)

High initial structure similarity is observed between Omicron-RBD and WT-RBD with the RMSD of 0.336. RMSF analysis showed that higher fluctuations were observed in the binding site hairpin loop regions especially between 478K to 485G(Figure 4B). Presence of disulphide bonds between (CYS-480 to CYS-488) is majorly responsible for omicron RBD folding and stability(Figure: 1F).

**Figure 4:**
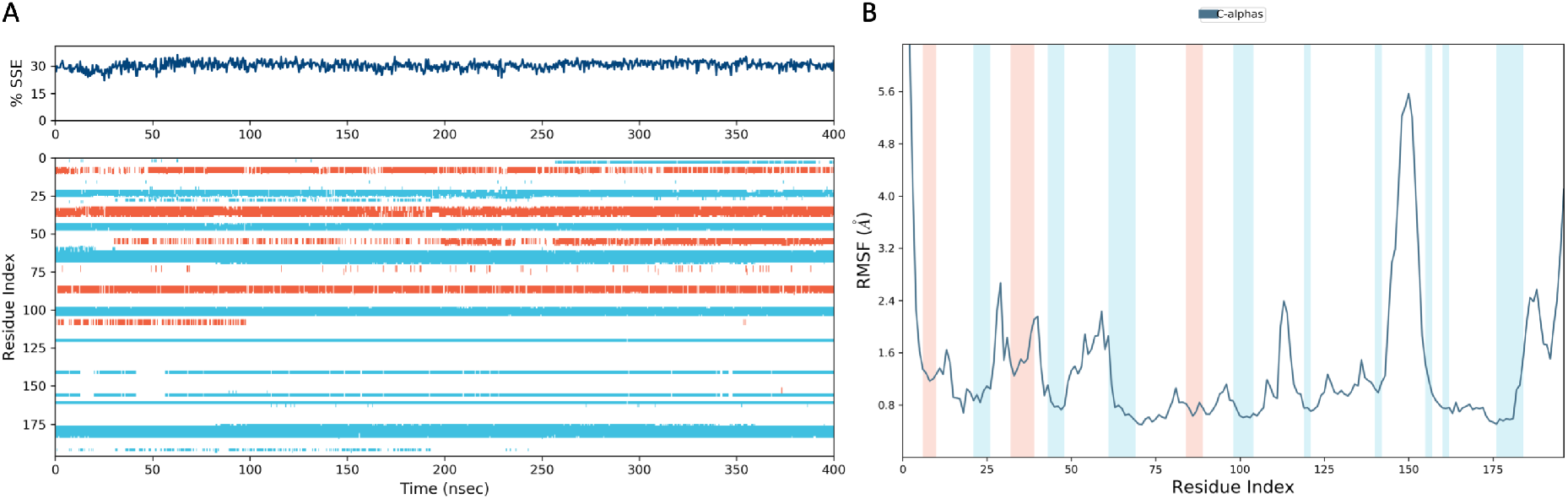
Secondary structure analysis over 400ns production run. **(A)** Secondary Structure Elements analysis of omicron-RBD **(B)** RMSF analysis of Omicron-RBD calculated during whole production run of 400ns.

Secondary structure elements analysis showed that Omicron RBD consists of 8.95% helix 21.50% Strand with 30.45% total SSE. SSE components on the whole trajectory showed that alpha helices region in between SER-438S to SER-443 slowly disappeared during the evolution of protein complex (Figure 4A).

Hydrogen bond formation between omicron RBD and h-ACE2 defines the structural stability of the protein complex. H-bond occupancy analysis is done by using VMD H-bond tool has provided major residues involved in omicron RBD and h-ACE2 binding interaction.

The h-ACE2 major residues involved in H-bond interactions are GLU35, TYR41, TYR83, GLN24, LYS353, ASP38, HIS34, whereas in omicron RBD ARG493, THR500, ASN487, TYR489, ALA475, GLY502, SER496,TYR453, GLY476 residues are involved in H-bond formation. THR500-TYR41(51.25%) and GLY502-LYS353(23.68%) residue combinations are majorly occupying H-bonds in the total production run of omicron RBD and h-ACE2 complex. Where as in Wild type RBD TYR489-TYR83, THR500-TYR41, TYR83-ASN87, GLY502-LYS353 are majorly occupying H-bonds. Unique H-bond interactions ASN-487-TYR83, SER496-ASP38, TYR453-HIS34, GLN24-GLY476 were observed between omicron RBD-h ACE2. (Figure 5).

**Figure 5:**
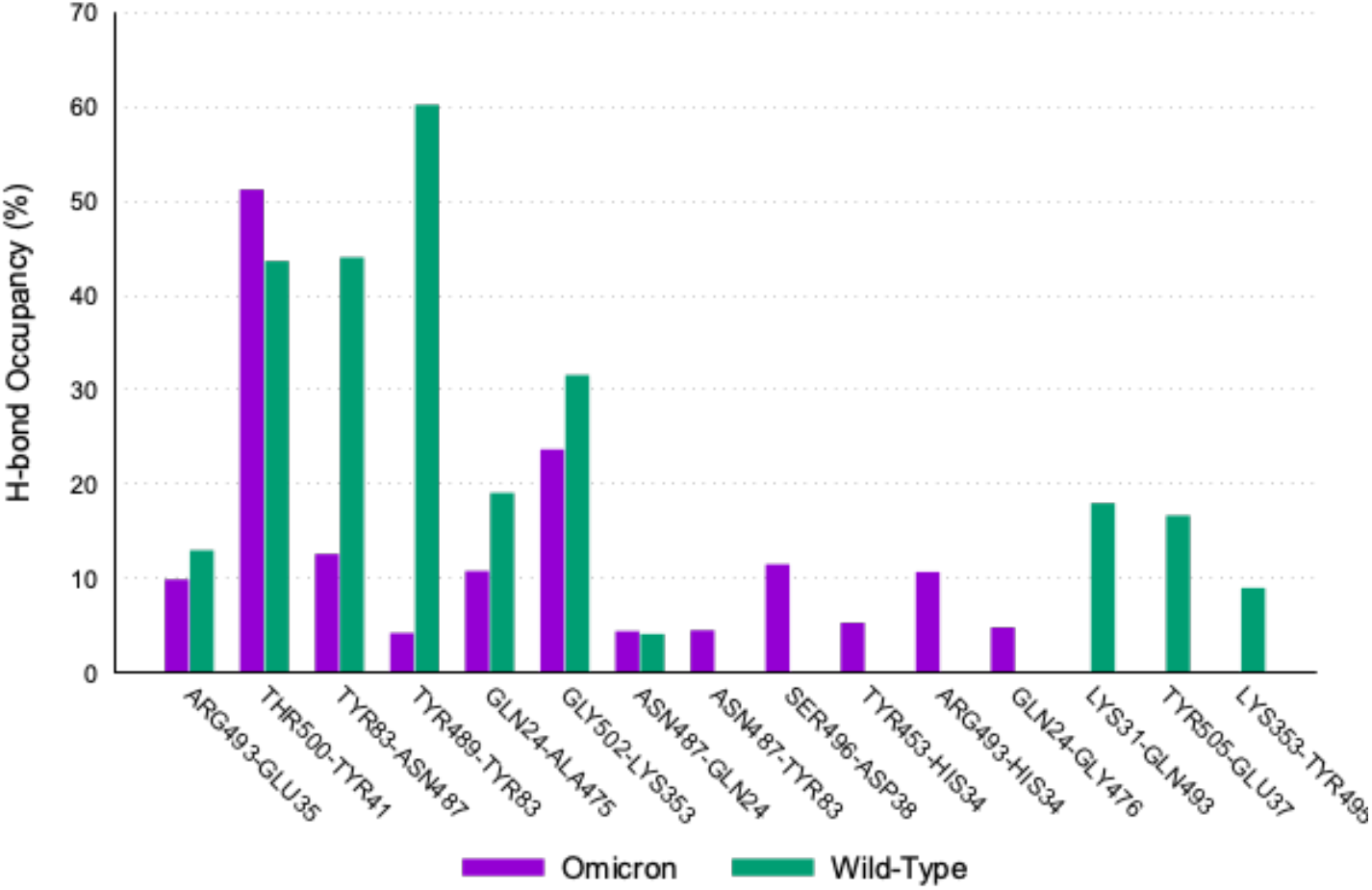
Comparative Hydrogen bond occupancy between RBD and h-ACE2 both in omicron and wild type during 400ns molecular dynamic simulations.

### Molecular Docking and Interaction Analysis

All the peptides with respective mutated residues and reference peptide were docked with omicron-RBD considering the interacting residues of peptide and protein obtained as a part of protein interacting analysis as the active site residues that coincided with literature data. The fourteen peptides which scored more than the reference peptide (−109.1±6.3kcal/mol) ranging between (−114.0±8.8 to −127.8±2.1kcal/mol) has provided binding and affinity insights on the core binding site. Peptides with highest scores are subjected to further analysis and are tabulated below (Table: 1). Respective amino acids that scored highest are replaced at 13 positions resulting in mutated peptides (mutated residues are showed in red colour).

**Table 1:**
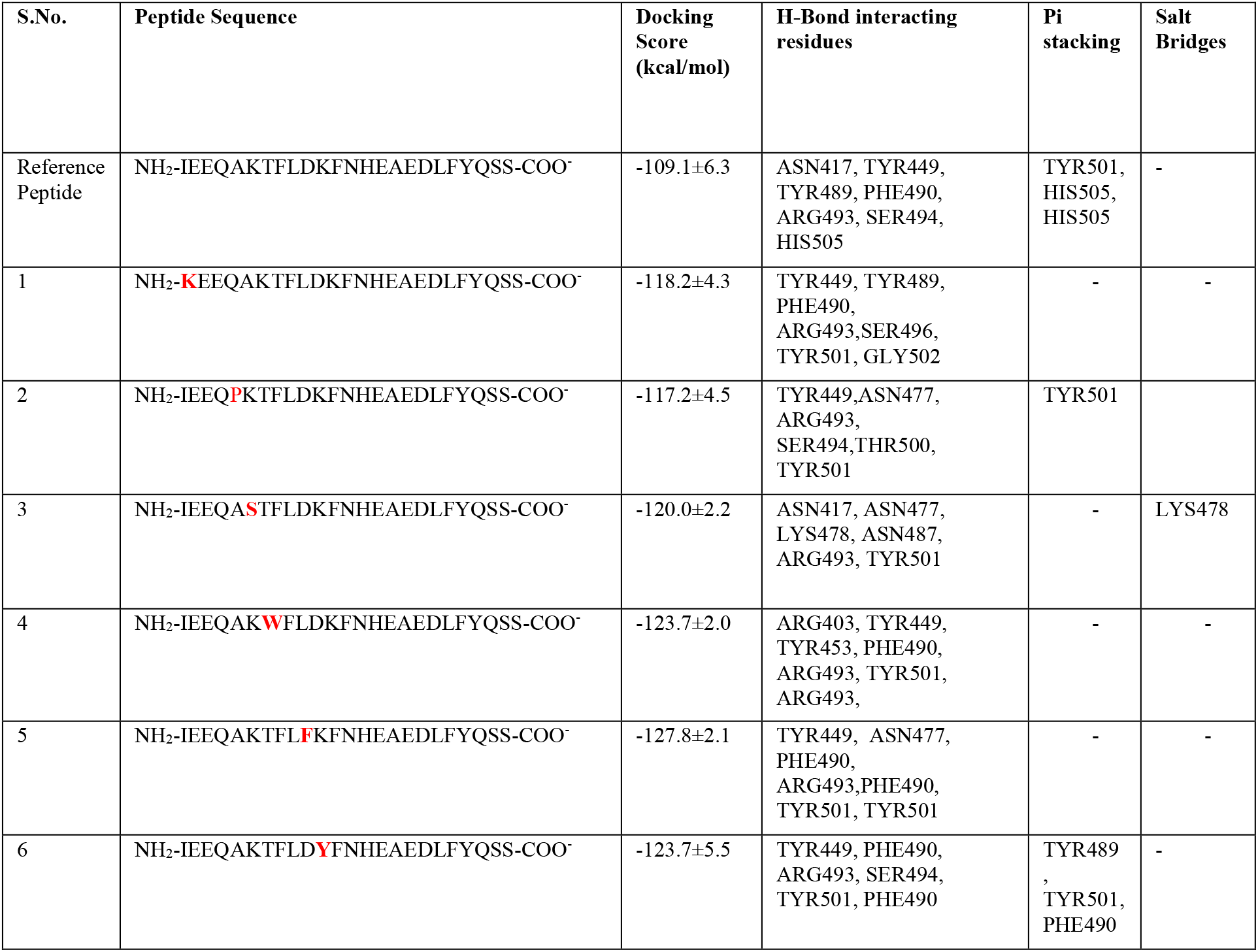
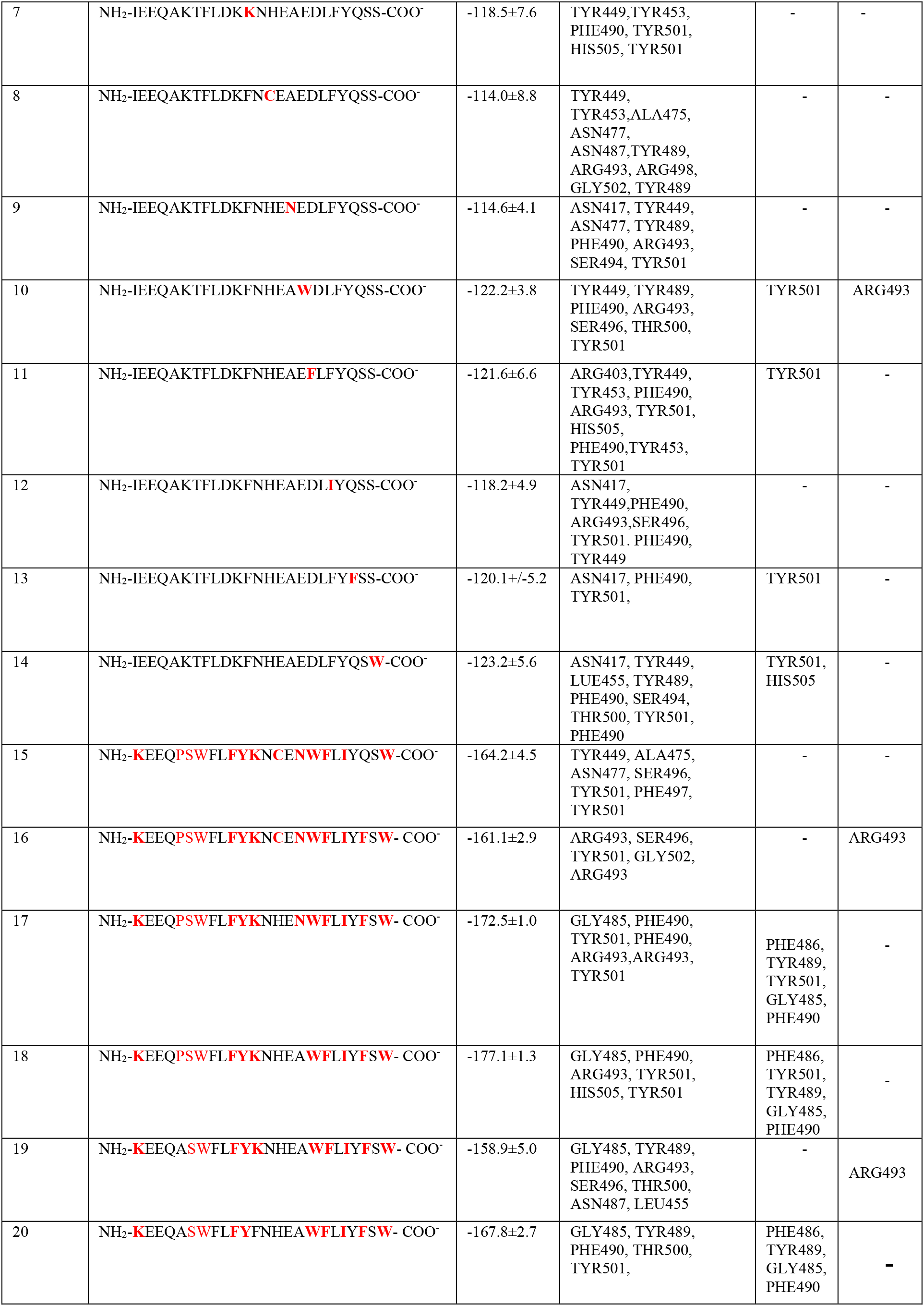
Reference h-ACE2 along with Biomimetic peptides with top binding scores and hydrogen bond interacting residues.

Comparatively the mutated peptides scored better than the reference from where clues are drawn for mutating the initial residues with high scored ones at respective positions resulting in peptides that scored thrice than reference.

**Figure 6:**
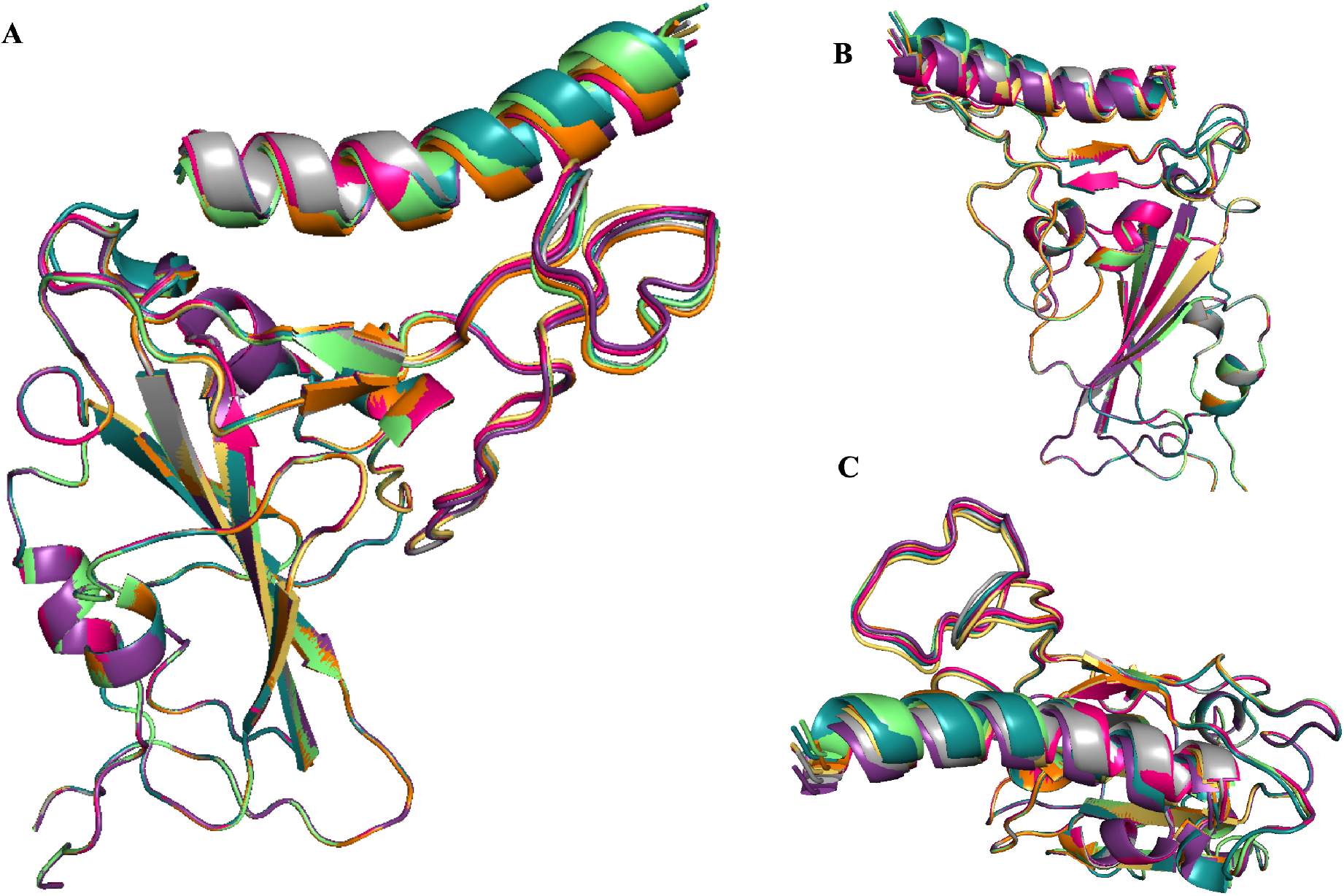
Superimposition of docked poses of Highly scored peptide mimetics along with wild type peptide. (wild type(peptide-1): Orange, peptide-15:Green, Peptide-16:Teal, Peptide-17:Pink, Peptide-18:Gold, Peptide-19:Violet, Peptide-20:Gray **(A)** Front View of RBD along with peptides **(B)** Back view of omicron RBD with Peptides **(C)** Top View of Omicron RBD with peptides

Replacement of uncharged nonpolar residues with positively charged amino acid resulted in the highest scored peptide with score of – 177.1±1.3 kcal/mol.

## Conclusion

Host receptor recognition and attachment by virion is facilitated by interface of omicron RBD with h-ACE-2. Thus, design of peptides that have resemblance with h-ACE2 can inhibit the protein-protein interaction and be a potential therapeutic in order to combat the virus. The modelled complex is subjected to molecular dynamic simulation for insights into the atomistic details of protein-protein interactions. Ile21 to Ser44 residues of h-ACE-2 are considered as key residues that interact with omicron RBD domain and are mutated to generate a library of 453 novel peptides. Out of this, thirteen peptides exhibited stronger interaction with RBD than that of ACE-2 which are considered to be worthy for further investigation. Based on the interaction analysis of 14 bio mimetic peptides design of novel peptides is done that demonstrate higher binding affinity than wild type h-ACE2 derived peptide.

## Acknowledgments

NIPER-Mohali computing facility is used for all computational studies. GNU plot is used to depict the statistical studies. Throughout the study process, Pymol is used to generate high quality images.

